# The impact of different sources of heterogeneity on loss of accuracy from genomic prediction models

**DOI:** 10.1101/374355

**Authors:** Yuqing Zhang, Christoph Bernau, Giovanni Parmigiani, Levi Waldron

## Abstract

Cross-study validation (CSV) of prediction models is an alternative to traditional cross-validation (CV) in domains where multiple comparable datasets are available. Although many studies have noted potential sources of heterogeneity in genomic studies, to our knowledge none have system atically investigated their intertwined impacts on prediction accuracy across studies. We employ a hybrid parametric/non-parametric bootstrap method to realistically simulate publicly available compendia of microarray, RNA-seq, and whole metagenome shotgun (WMS) microbiome studies of health outcomes. Three types of heterogeneity between studies are manipulated and studied: imbalances in the prevalence of clinical and pathological covariates, 2) differences in gene covariance that could be caused by batch, platform, or tumor purity effects, and 3) differences in the “true” model that associates gene expression and clinical factors to outcome. We assess model accuracy while altering these factors. Lower accuracy is seen in CSV than in CV. Surprisingly, heterogeneity in known clinical covariates and differences in gene covariance structure have very limited contributions in the loss of accuracy when validating in new studies. However, forcing identical generative models greatly reduces the within/across study difference. These results, observed consistently for multiple disease outcomes and omics platforms, suggest that the most easily identifiable sources of study heterogeneity are not necessarily the primary ones that undermine the ability to accurately replicate the accuracy of omics prediction models in new studies. Unidentified heterogeneity, such as could arise from unmeasured confounding, may be more important.

## 1. Introduction

Quantification of heterogeneity between studies and its impact on validation of decision models is important across a wide range of applications. It has been noted that independent validation of genomic classifiers is rare (Castaldi *and others*, 2011), and the difficulty of external validation and study heterogeneity is common not only in microarray studies but in GWAS (König, 2011), and RNA-seq studies (Xu *and others*, 2016). External validation is critical in any research domain affected by heterogeneous samples, sample selection bias, or technical batch effects. However, it has proven especially difficult for classifiers and subtypes identified from gene expression data. Patient populations can be heterogeneous in their exposures, geography, race/ethnicity, and socioeconomic status, and these differences could manifest as biologically distinct forms of diseases that vary systematically between studies. Batch effects (Leek *and others*, 2010) and platform effects impact on reproducibility across studies. The sources of batch variation, sampling bias, and other heterogeneity may be unknown. Finally, “samples of convenience” are the norm in translational genomics research (Simon *and others*), due to the difficulty of collecting tissue specimens and the time-consuming process of clinical follow-up. In spite of these challenges, we expect clinically-relevant genomic findings to be reproducible at hospitals with different populations around the world, suggesting robustness in the presence of some heterogeneity. The “molecular portraits” of breast cancer, for example, have been broadly reproduced across platforms and centers (Hu *and others*).

Previous studies have shown that accuracy estimates of genomic prediction models based on independent validation are inferior to cross-validation estimates (Castaldi *and others*, 2011; Bernau *and others*, 2014), but did not identify the sources of heterogeneity responsible. Ma *and others* (2014) and Chang and Geman (2015) showed, by learning diagnostic models on large number of studies, how to estimate whether enough heterogeneity has been explored to achieve a desired degree of generalizability. Specifically, Chang and Geman (2015) showed that cross-study validation (CSV) error rate exceeds the randomized cross-validation (RCV) error rate for any number of studies. The latter increases with the diversity of studies, and both converge to the optimal rate for the whole population. Methods for correction of validation accuracy estimation in training samples of biased covariate distribution to the unbiased distribution (Cortes *and others*, 2008) and to the covariate distribution in test samples (Uno and Inoue, 2017) have been proposed.

It remains critical to understand the contributions of each component of “study effects” to the difference of performance. In practice, a standard approach is to remove as many sources of heterogeneity as possible, such as Waldron *and others* (2014) and Riester *and others* (2014), which limited meta-analyses to late-stage, high-grade, serous ovarian cancer. Similarly, recommendations for the replication of genome-wide association studies include studying a “similar” population. However in many cases it is unclear what measures of study similarity are important, and unnecessarily restrictive inclusion criteria have costs in reduced sample size and loss of generality of findings. Thus the question arises of which sources of heterogeneity do in fact impact the accuracy of cross-study prediction, and how these can be determined from the data. We propose that the impact even of still-unidentified sources of heterogeneity can be accounted for and quantified if independent studies are available.

We compare within and across study validation of omics-based prediction models using simulations which are generated from publicly available datasets, including both microarray and sequencing data. We investigate the impact of three possible types of heterogeneity on cross-study validation performance: changes in prevalence of known clinical and pathologic factors, changes in gene expression covariance structure for example due to batch or platform effects, and changes in the true models associating gene expression and clinical factors with outcome. These sources of heterogeneity are manipulated and equalized in turn, while we compare within-to across-study validation of predictions for the disease outcome or risk scores for survival. The methodology of this study can be applied to investigating the effects of study heterogeneity on model validation in any scenario where multiple independent but comparable datasets are available.

## 2. Methods

We evaluate the effects of across-study heterogeneity by resampling of studies, and preserving the distribution and covariance of gene expression through resampling of individuals within studies. We generate linear models associating clinical/pathological variables and gene expression to the outcome based on the original experimental data. We use these models to generate new outcome variables, which preserve the properties of the original outcome. We emphasize that standard clinical factors were included as required, unpenalized covariates in the “true” prognostic model so that their associations with outcome would be guaranteed to be preserved across simulations.

We review the simulation procedure in the following sections, which involves a 3-step bootstrap method. To implement this simulation approach, we developed the *simulatorZ* package and made this available through Bioconductor. Scripts for reproducing the results of this paper are stored and documented at https://bitbucket.org/zhangyuqing/datasetheterogeneity.

### 2.1 Datasets

#### 2.1.1 Microarray studies

Public experimental studies are hereafter referred as the “original” datasets. Steps for preprocessing these datasets are detailed in Supplementary Materials.

The first compendium consists of 7 curated breast cancer datasets, which was originally published by Haibe-Kains *and others* (2012), and used in Bernau *and others* (2014). These studies contain censored disease and metastasis-free survival (DMFS) response, and 1021 estrogen-receptor positive breast cancer individuals. Available covariates in these studies include patient age at diagnosis, tumor size, and histological grade. Age and tumor size were dichotomized at 50 years and 2cm, respectively (Haibe-Kains *and others*, 2012), and grade was kept at three levels (low/medium/high) as in the original datasets. Synthesized Hazard Ratios (HR) of the tumor size, grade and patient age are 1.96, 1.65 and 1.2 (Supplementary Table 2). This is consistent with commonly used prognostic factors for primary breast cancer such as the Nottingham Prognostic Index (Haybittle *and others*, 1982), showing that the dichotomized tumor size and grade are prognostic. Sample information and distributions of covariates are summarized in Table 1.

**Table 1.** Covariate distributions in the ER-positive breast cancer microarray datasets. Percentages are rounded to the nearest tenth. Datasets acronyms: CAL = University of California, San Francisco and the California Pacific Medical Center (United States), MNZ = Mainz hospital (Germany), TAMl and TAM2 represents SUPERT AM_HGU133A and SUPERT AM_HGU133PLUS2, which are provided by Haibe-Kains *and others* (2012), TRB = TransBIG consortium dataset (Europe), UNT = the cohort of untreated patients from the Oxford Radcliffe Hospital (United Kingdom), VDX = Veridex (the Netherlands). Column labels: #patients = the number of patients after cleaning. %> 2cm = percentage of patients in cleaned datasets with tumor size larger than 2cm. %low, medium and high-grade = percentage of patients with low, intermediate and high level of histological grade. %> 50yrs = percentage of patients older than 50 years.

| NO. | Name | #patients | %>2cm | %high-grade | %medium-grade | %low-grade | %>50yrs |
| --- | --- | --- | --- | --- | --- | --- | --- |
| 1 | CAL | 68 | 63.2 | 57.4 | 35.3 | 7.4 | 61.8 |
| 2 | MNZ | 162 | 40.7 | 9.9 | 73.5 | 16.7 | 76.5 |
| 3 | TAM1 | 317 | 58.0 | 18.3 | 58.4 | 23.3 | 91.2 |
| 4 | TAM2 | 128 | 53.1 | 31.3 | 44.5 | 24.2 | 93.0 |
| 5 | TRB | 132 | 43.2 | 26.5 | 51.5 | 22.0 | 29.5 |
| 6 | UNT | 72 | 38.9 | 16.7 | 43.1 | 40.3 | 59.7 |
| 7 | VDX | 142 | 2.8 | 71.8 | 25.4 | 2.8 | 57.7 |
|  | overall | 1021 | 44.1 | 29.6 | 50.9 | 19.5 | 72.3 |

The other collection, of 5 microarray datasets containing 935 ovarian cancer patients, was published in Ganzfried *and others* (2013) and is available from the *curatedOvarianData* Bioconductor package. Patients with late-stage, high-grade cancer were included. The datasets contain patient age and debulking status as available covariates. Age was dichotomized at 70 years. Synthesized Hazard Ratios of these covariates are 1.84 (age) and 1.48 (debulking), as shown in Supplementary Table 3. Distributions of these covariates are summarized in Supplementary Table 4.

#### 2.1.2 The RNA-seq study

In addition to the microarray studies, we created simulations in RNA-seq data with time-to-event outcome. Lacking sufficient RNA-seq studies for a full cross-study validation, we substituted the TCGA microarray dataset with TCGA RNA-seq data in the ovarian cancer study collection. The TCGA RNA-seq study is also available in the *curatedOvarianData* package. It contains 190 patients, a subset of the microarray cohort. We used the same clinical covariates, patient age and debulking status, for the simulations using the RNA-seq study.

#### 2.1.3 Metagenomics studies

We further evaluated the impact of heterogeneity in whole-metagenome shotgun-sequencing studies with binary disease outcomes, which are available from the *curated-MetagenomicData* Bioconductor package (Pasolli *and others*, 2017). We focused on 3 studies with gut microbiome samples of type-II diabetes (T2D) patients and healthy individuals. The binary outcome indicates the disease status for each sample. We identified Body Mass Index (BMI) as the covariate to be balanced for these studies.

For predictors we used gene families, as estimated by HUMAnN2 (Abubucker *and others*, 2012) and provided by *curatedMetagenomicData*, in the prediction models. We performed a series of feature selection steps, as documented in Supplementary Materials, after which 800 features were included in the studies. We dichotomized BMI at 25 based on the empirical separation of BMI as “under to normal weight” and “overweight to obese”. Supplementary Table 5 and Figure 2 summarize the sample and covariate information of these studies.

### 2.2 Simulation approach and the simulatorZ Bioconductor package

Bernau *and others* (2014) introduced a systematic approach for synthesizing a group of independent microarray datasets with survival outcome for cross-study assessment of prediction methods. We developed the *simulatorZ* package to create collections of independent genomic datasets with realistic properties and outcome variables generated from a known risk model. *simulatorZ* also implements the Màs-o-menos algorithm (Zhao *and others*, 2014; Donoho and Jin, 2008) and provides basic facilities for cross-validation and cross-study validation of prognostic models.

#### 2.2.1 Simulation of independent datasets

The simulation procedure contains three steps. The first is a non-parametric bootstrap at the dataset level, in which studies are sampled with replacement from the list of original studies. This estimates the variability due to sampling of studies from a “super-population” of studies (Hartley and Sielken, 1975). The second step is another non-parametric bootstrap at the patient level, where observations are sampled with replacement from each dataset selected in step 1. In the final step, a linear model is fit to the *original* datasets, then used to simulate new outcome on the simulated datasets (parametric bootstrap).

For studies with time-to-event outcome, we fit a proportional hazard (PH) model to the data:

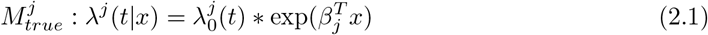

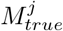 is the PH model for the *j*-th dataset, whose hazard function is λ^*j*^(*t|x*), with *x* as covariates. 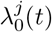 is the baseline hazard function for this study. *β* represents the regression coefficients.

This generative model in step 3 combines the truncated inversion method of Bender *and others* (2005), the Nelson-Aalen estimator (Nelson, 1969, 1972; Aalen, 1978) for cumulative hazard functions, and the *CoxBoost* method of generating best-fit linear risk scores (Binder and Schumacher, 2008). We first use *CoxBoost* to obtain coefficients of linear predictors fitted to the original datasets, using the genes plus the clinical covariates, such as tumor size, debulking, histological grade and patient age, as predictors. The prognostic covariates were included to be mandatory unpenalized. We also obtained the Nelson-Aalen estimator of baseline cumulative survival and censoring hazard, which, together with the *CoxBoost* coefficients and Equation 2.1, define the “true” models of survival for each dataset. Finally, we have:

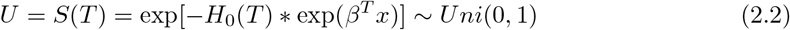

*H*_0_ is the baseline cumulative hazard for the lifetime random variable *T*. *S* denotes the survival function. We sample two independent, uniformly distributed variables u_1_ and u_2_, then simulate the survival (*T*) and censoring (*C*) time, with

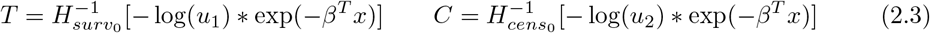

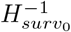 and 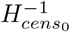 are the inverses of baseline cumulative survival and censoring hazard, respectively. These are inverted by finding the point on the time line such that the values calculated by *−* log(*u*) *\**exp(*−β^T^ x*) are closest in absolute value to the cumulative hazards. The simulated survival response is the smaller one between *T* and *C*.

For studies with binary outcome, we fit a logistic regression to the data:

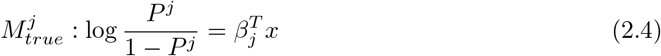

*P^j^* denotes the probability of a sample in study *j* to have the disease. Here, for studies with binary outcome, the term “true model” refers to Equation 2.4 together with the model coefficients *β*. After fitting the model and estimating the probability 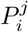 for a sample *i* in study *j*, we draw a value from the Bernoulli distribution 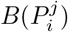 as the simulated outcome for that sample.

#### 2.2.2 Summary of within and across-study model performances

We selected the Más-o-menos algorithm (Zhao *and others*, 2014; Donoho and Jin, 2008) and ridge regression (Hoerl and Kennard, 1970) as examples of predictive models to generate risk scores on the simulated datasets. Bernau *and others* (2014) and Zhao *and others* (2014) have shown that these algorithms perform comparably to more complicated methods in the microarray datasets that we use. We repeated 100 simulations and model validations for each dataset/algorithm combinations. These include applying Más-o-menos and ridge regression to each collection of datasets, including breast and ovarian cancer microarray studies, ovarian cancer microarray and RNA-seq compendium, and the metagenomic studies. We estimated the model accuracy with C-indices for studies with time-to-event outcome, and area under ROC curve (AUC) for studies with binary outcome.

In each of 100 iterations, we simulated a list of independent datasets of sample size *n* = 150 using the “original studies” and the 3-step bootstrap approach. We then generated a matrix of accuracy estimates for all combinations of training and test sets, as described by Bernau *and others* (2014). Cross-study validation (CSV) performance was summarized by the simple average of accuracy estimates for training and validation across all pairs of independent studies (off-diagonal elements of the matrix), and performance of the 4-fold cross-validation (CV) was summarized by the average of the diagonal elements (Bernau *and others*, 2014). The process was repeated while altering potential sources of across-study heterogeneity, as described in Sections through 2.5. The above methods are summarized in Figure 1.

**Fig. 1.**
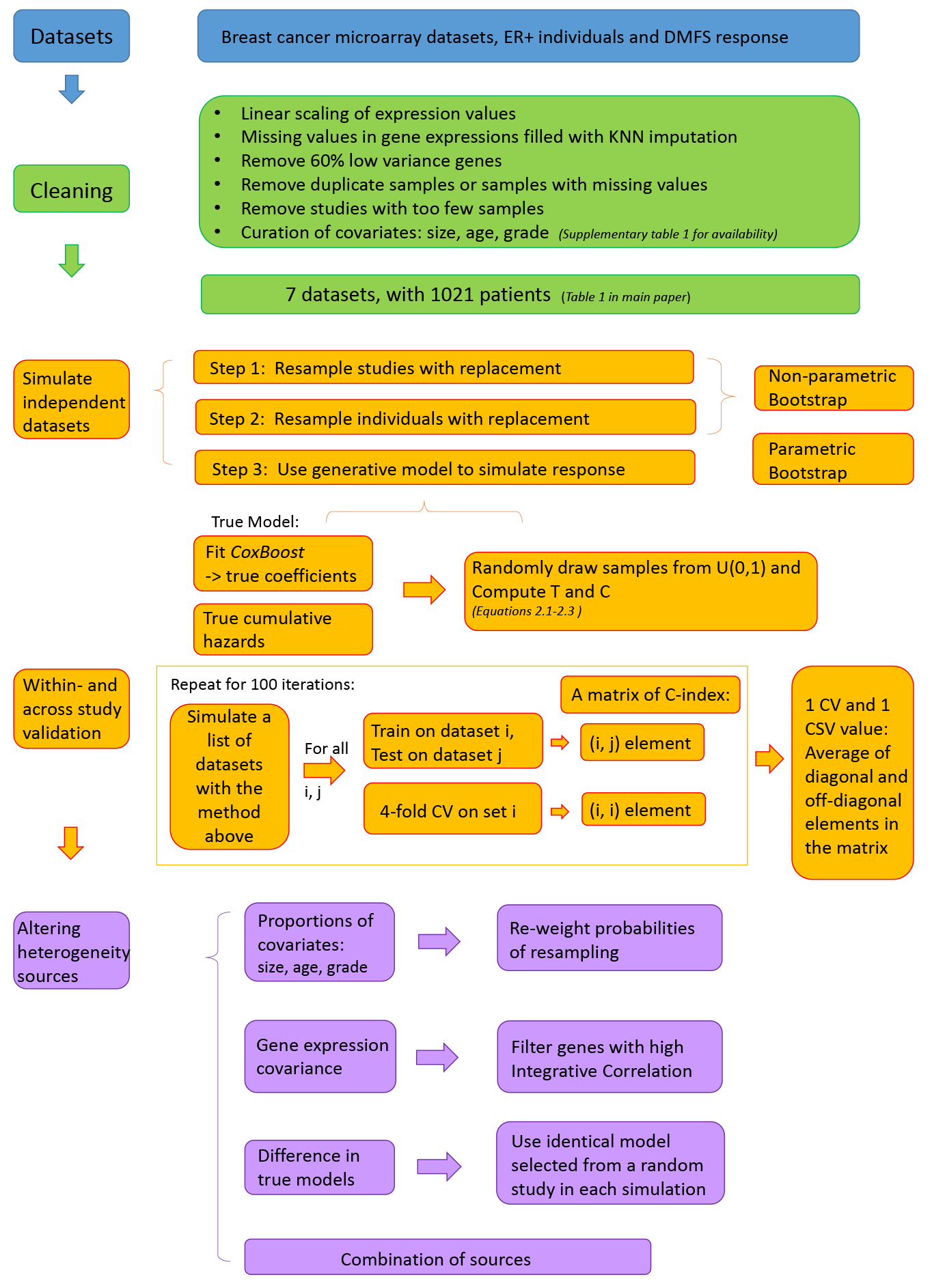
A schema of our study. Simulation methods (using the breast cancer microarray studies as an example) are summarized in this flow chart.

#### 2.2.3 Comparison of model accuracy using the RNA-seq study

The methods for the RNA-seq study differed due to the availability of only a single RNA-seq study with comparable outcome to the microarray studies. We compared cross-validation within this study to training and validation across different data types (one microarray and one RNA-seq study). During the simulations, we skipped the first bootstrap step which re-samples studies from the collection, in order to keep track of the RNA-seq study among the other microarray datasets. We generated the matrix of accuracy estimates as before. When comparing model performances between within and across-study validation, we limited the comparison to only when the RNA-seq study is involved. This means that, in each simulation, we select the row and the column of the performance matrix which correspond to the RNA-seq study. The performance of cross-validation is the diagonal element located at the intersection of the row and the column. The cross-study performance is summarized by averaging over all remaining elements within the row and the column.

### 2.3. Clinical covariates

To simulate studies with similar distributions of a single clinical covariate, such as dichotomized tumor size, debulking, or of young and old patients, we first define the overall proportions of patients for that covariate using the proportions in the union of all original studies. We then changed the probabilities for bootstrap resampling on the individuals, so that on average, each simulated study would have proportions of patients equal to the overall proportions of the covariate. To balance on multiple covariates, we re-weighted individual bootstrap sampling probabilities of each patient to result in identical joint probability distributions of the clinical/pathologic covariates across datasets. For example, we denote the proportion of individuals with age *>* 70*yrs* and suboptimal debulking status in all ovarian cancer studies combined as *P_common_*. Such proportion in a study *s* alone is denoted as *P_s_*. Then in the simulated study resampled from study *s*, patients with age *>* 70*yrs* and suboptimal debulking status will be resampled with a probability that is proportional to *P_common_/P_s_*, which is scaled to one for all individuals in study *s*. Re-weighting of the sampling probabilities, rather than enforcing strict equality of proportions, reflects the reality that these proportions are subject to sampling variation.

#### 2.3.1 Mixed-effect models

We quantify the impact of changing the proportion of covariates on the cross-study prediction accuracy by a regression approach, implemented via mixed-effect models. These are implemented in the two compendia of microarray studies. We established a “baseline” scenario of 100 simulations based on the original studies, without any manipulation of sources of heterogeneity. We then performed another 100 simulations where we changed the bootstrap re-sampling probabilities at the patient level to produce, on average, equal distributions of covariates in each study. This scenario is called “balancing covariates”. These simulations produce 100 matrices of C-index for each of the two scenarios, “baseline” and “balancing covariates”.

The validation matrices from the two scenarios are then paired together: the first matrix from “baseline” is paired with the first one from “balancing covariates”, then the second is paired with the second,…,the 100th is paired with the 100th. These pairs of validation matrices, as well as their corresponding lists of simulated independent datasets, are used to compute the changes in cross-study validation accuracy, and the changes in the proportions of subjects in each covariate subgroup. For each off-diagonal position in the matrix, we calculate the change in C-index as the value in “balancing covariates” at that position minus the corresponding value in “baseline”. Also, for the same position, there is a pair of training/test sets for “baseline”, and another for “balancing covariates”. We compute the difference between the proportions of patients in the training (test) set for “balancing covariates” and the proportions in the training (test) set for “baseline”. This difference will serve as a potential predictor for changes in the C-index.

We then fit the model:

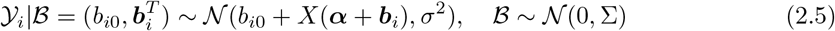

For *n* independent studies in a single simulation *i*, *Y_i_* represents the changes in C-index. Each *Y_i_, i* = 1, 2, …, 100, is a vector of length *n* * (*n −* 1) corresponding to all possible training/test set pairs in the *i*-th simulation. The design matrix *X* contains predictors which include the changes in the covariate distributions in the training and the test set, as well as a third interaction term. Entries in *X* are grouped by simulation indices (*i*). *X* has *n* *(*n −* 1) *100 rows, accounting for all cross-study training/test set pairs across the 100 simulations. ***α*** = (*α*1, *α*2, *α*3)^*T*^ is a vector of fixed effects. The three dimensions correspond to the changes of covariate proportions in the training and test set, and the interaction term. In the *i*-th simulation, 𝓑 equals to 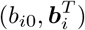, ***b***_*i*_ = (*b_i_*_1_*, b_i_*_2_*, b_i_*_3_)^*T*^. They serve as a vector of normally distributed random effects, with mean and variance estimated across the only level of grouping, 100 simulations.

Having the coefficients vary across simulations accounts for the potential inherent correlations in the observations within each simulation group, which uses the same list of simulated datasets. The observations are independent across groups, given that the simulated datasets are independent. It is modeled as having no intercept in the fixed component because the average difference over many simulations must be zero when there is no difference in the covariate distribution or the generating model for these two validation matrices. However there will be random variation across simulations, for which we include an intercept in the random component. This model is fit on every covariate for the *microarray* dataset/algorithm combinations.

### 2.4 Expression covariance

To investigate the potential impact of heterogeneity between gene expression levels in different datasets, we compared the baseline case to the case where we only use genes with high Integrative Correlation (Parmigiani *and others*, 2004; Garrett-Mayer *and others*, 2008) between every dataset pair. Briefly, we first calculated the Pearson correlation matrix of each gene expression matrix. For each pair of datasets, the Pearson correlation of the *k*-th rows of the two correlation matrices is the Integrative Correlation of gene *k*. We did a grid search for the threshold of the Integrative Correlation, such that around 1000 genes with the highest Integrative Correlation scores between every pair of *original* datasets were included. We also used arbitrary cut-offs 0.4 for breast cancer microarray studies and 0.2 for ovarian cancer microarray datasets, as a comparison with selecting 1000 genes. Simulations in the microbiome studies are also performed with arbitrary cut-offs 0.4.

### 2.5 True models

We equalized the “true models” of each dataset. In each simulation, we randomly select an original study in the 3rd bootstrap step, and use its corresponding “true model” to simulate the new outcome for all re-sampled studies from the first two bootstrap steps. For studies with time-to-event outcome, we equalized separately the coefficients of the linear risk score, and the baseline hazard function. For studies with binary outcome, we used identical model coefficients.

## 3. Results

We first detail our observations from the cancer microarray studies of patient survival because they provide the greatest sample size, then summarize RNA and whole-metagenome sequencing results which are broadly consistent with these results. In original and simulated data, we observed a substantial loss of prediction accuracy in cross-study validation (CSV) when compared to cross-validation (CV). Reductions were approximately 0.04 on the C-index scale, and 0.12 on the AUC scale. We manipulated aspects of the simulated data to establish that reducing heterogeneity in the sources we investigate is not sufficient to eliminate the gap between across- and within-study prediction accuracy. Using identical “true” models is most effective in reducing this difference. Figures 2 and 3 summarize our results across multiple omics platforms, showing that the three factors of interest cannot fully explain the loss of accuracy in independent validation.

**Fig. 2.**
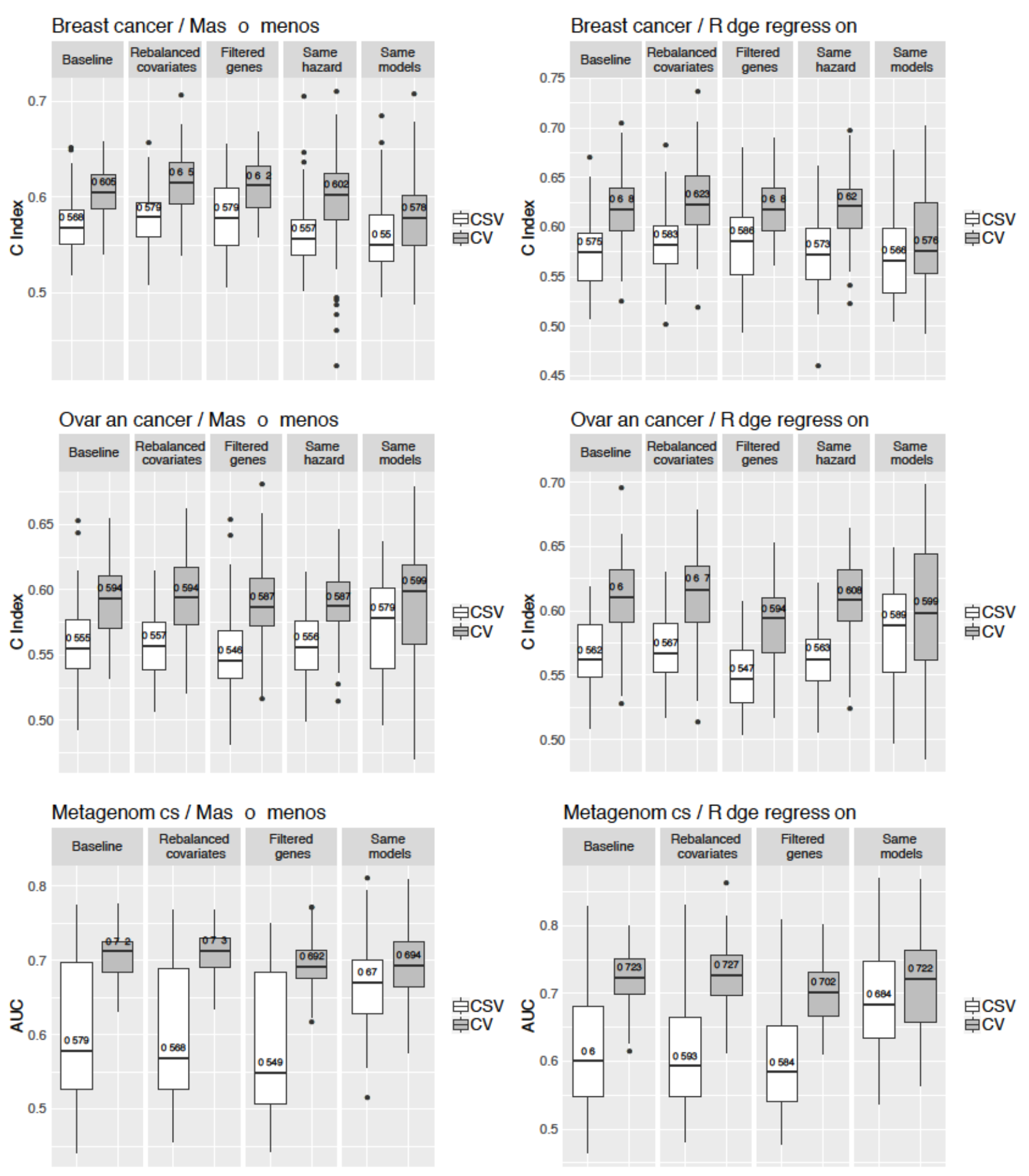
Simulation results comparing performances of cross-validation and cross-study validation. The "Baseline" scenario does not modify any source of heterogeneity. In "Re balanced covariates", we change the re-sampling probability to match the distribution of covariates. "Filtered genes" considers only genes with high Integrative Correlation. "Same hazard" uses the same cumulative hazard but different coefficients for simulations using microarray studies with time-to-event outcome. "Same models" uses the same data generating models. The numbers within the boxes show the median of the distributions. We observe that the sources of heterogeneity investigated in this work do not fully account for the loss of accuracy comparing across-to within-study validation.

**Fig. 3.**
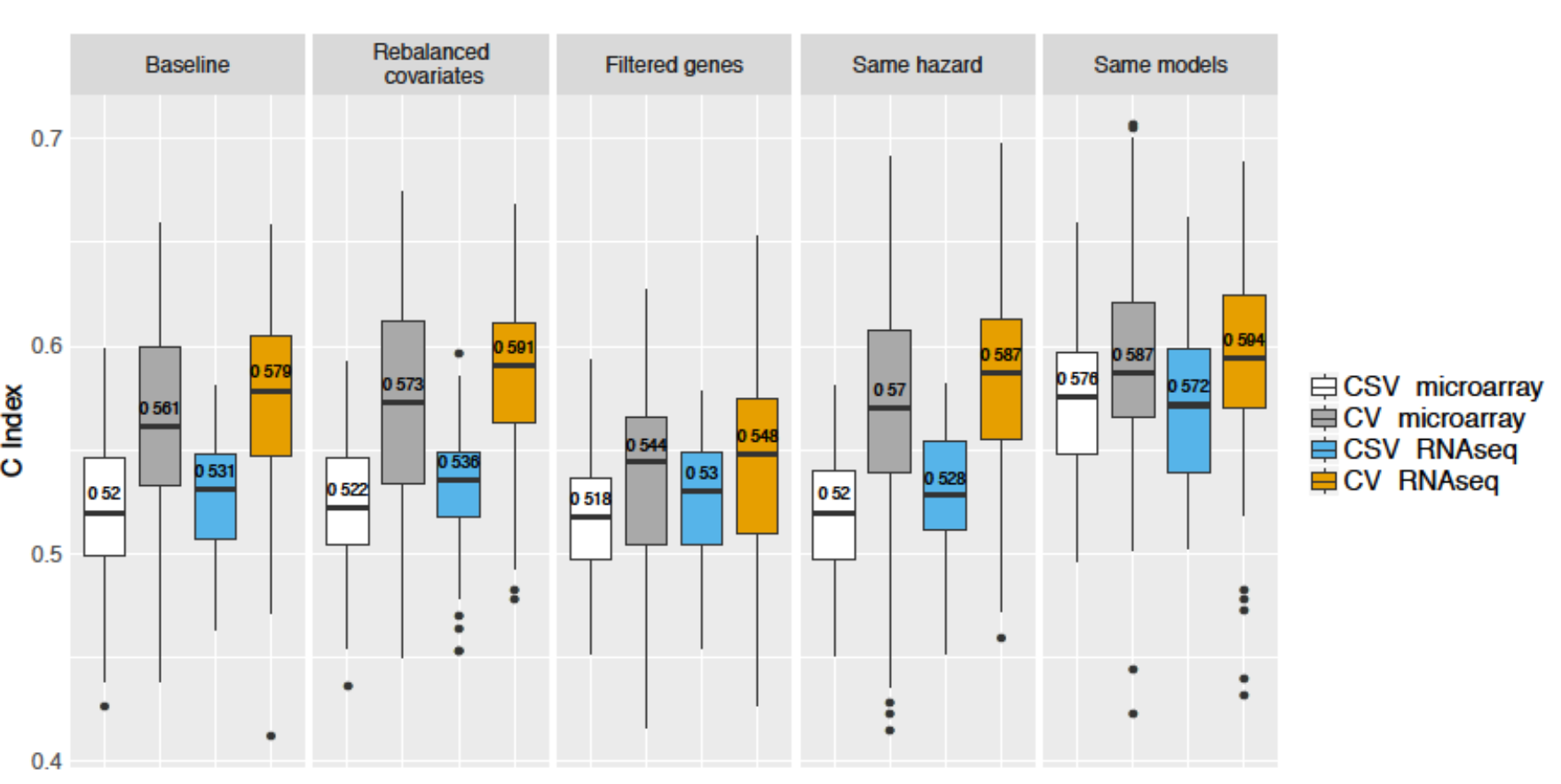
Simulation results comparing CSV to CV for evaluating Mas-o-menos risk prediction models in ovarian cancer microarray and RNA-seq studies. Colored boxes represent C-indices collected in within and across-study validation when RNA-seq data is involved.White/grey boxes represent results using only microarray studies. Note tha t the microarray simulations in this figure are *different from* those in Figure 2. In this figure, the comparison is limited to the TCGA study, which contains only the 190 overlapping samples between the microarray and RNA-seq platforms.

### 3.1 Simulation of microarray studies

Sections 3.1 through 3.4 are discussed in the context of the microarray studies. We re-capitulated the major properties of these two collections of studies in a 3-step bootstrap simulation procedure. The simulation resulted in within and across-dataset training validation characteristics comparable to those from prior clinical studies (Supplementary Figure 3). Most importantly, the simulation studies maintained a realistic difference in validation accuracy as estimated within and across studies by the C-index. This difference is seen for both ridge regression and the Más-o-menos method, in breast cancer and ovarian cancer (see panels “Baseline” in Figure 2).

### 3.2 Eliminating heterogeneity in clinical factors

To establish whether eliminating heterogeneity in known clinical factors could improve cross-study validation accuracy, we re-weighted bootstrap sampling probabilities to balance tumor size, grade and patient age for the breast cancer data, and age and debulking status for the ovarian cancer data. Proportions of these factors were then the same, in expectation, for each simulated dataset. For both cancer types, the differences between CV and CSV are not eliminated (Figure 2, “Rebalanced covariates”). In this scenario, differing distributions of these covariates does not contribute to the loss of prediction accuracy across studies relative to within studies.

Though the re-balancing of covariates does not mitigate the overall drop in accuracy in cross-study validation, we found that the rank of CSV scores may be somewhat affected depending on the algorithm used. Supplementary Figure 5 displays re-ordering of performance ranks for different training test pairs, but with no overall effect across all training testing pairs. To quantify this observation, we built a linear model for every covariate. Each model associates the changes in proportions of clinical factors to the changes in the CSV scores. Supplementary Table 6 summarizes the results of this analysis and highlights which covariates have an effect on CSV in the two methodologies used. Interestingly, even covariates that are prognostic of the survival outcome do not necessarily significantly relate to the prediction accuracy changes.

### 3.3 Filtering genes by Integrative Correlation

We filtered genes to include roughly the top 1000 genes with the highest Integrative Correlation (Parmigiani *and others*, 2004; Garrett-Mayer *and others*, 2008) between every pair of original datasets. We implemented this by searching on a grid for the required threshold of IC. The threshold is 0.24 for breast cancer (999 genes), and 0.15 for ovarian cancer (1002 genes). After filtering these genes on the original datasets, the differences between CV and CSV are only slightly reduced for breast cancer, and not reduced for ovarian cancer (Figure 2, “Filtered genes”).

As an alternative to fixing the size of the gene set, we used thresholds of 0.4 for breast cancer and 0.2 for ovarian cancer. Both CV and CSV perform slightly worse using the fixed thresholds compared to the grid search, as the stricter filtering results in loss of good predictors. The observed pattern remains that the gap in accuracy is not eliminated (Supplementary Figure 6), which suggests that the result is robust to the choice of thresholds. Filtering genes to enforce similar covariance structures across studies, as would be expected in the absence of substantial microarray batch or platform effects, does not remove the CV-CSV difference.

### 3.4 Using the same true model

True models of the experimental sets with time-to-event outcome differ in both baseline survival and risk coefficients. Figure 4 shows the average probability of survival in each dataset and for all datasets combined, which explains what we mean by the differences in the true models. Differences in baseline survival mean that survival across all patients is better in some datasets than others, for example varying between 60% and nearly 90% 5-year survival. We equalized both coefficients and baseline hazards across studies, while allowing the joint distribution of covariates and the gene covariance structure to remain heterogeneous.

**Fig. 4.**
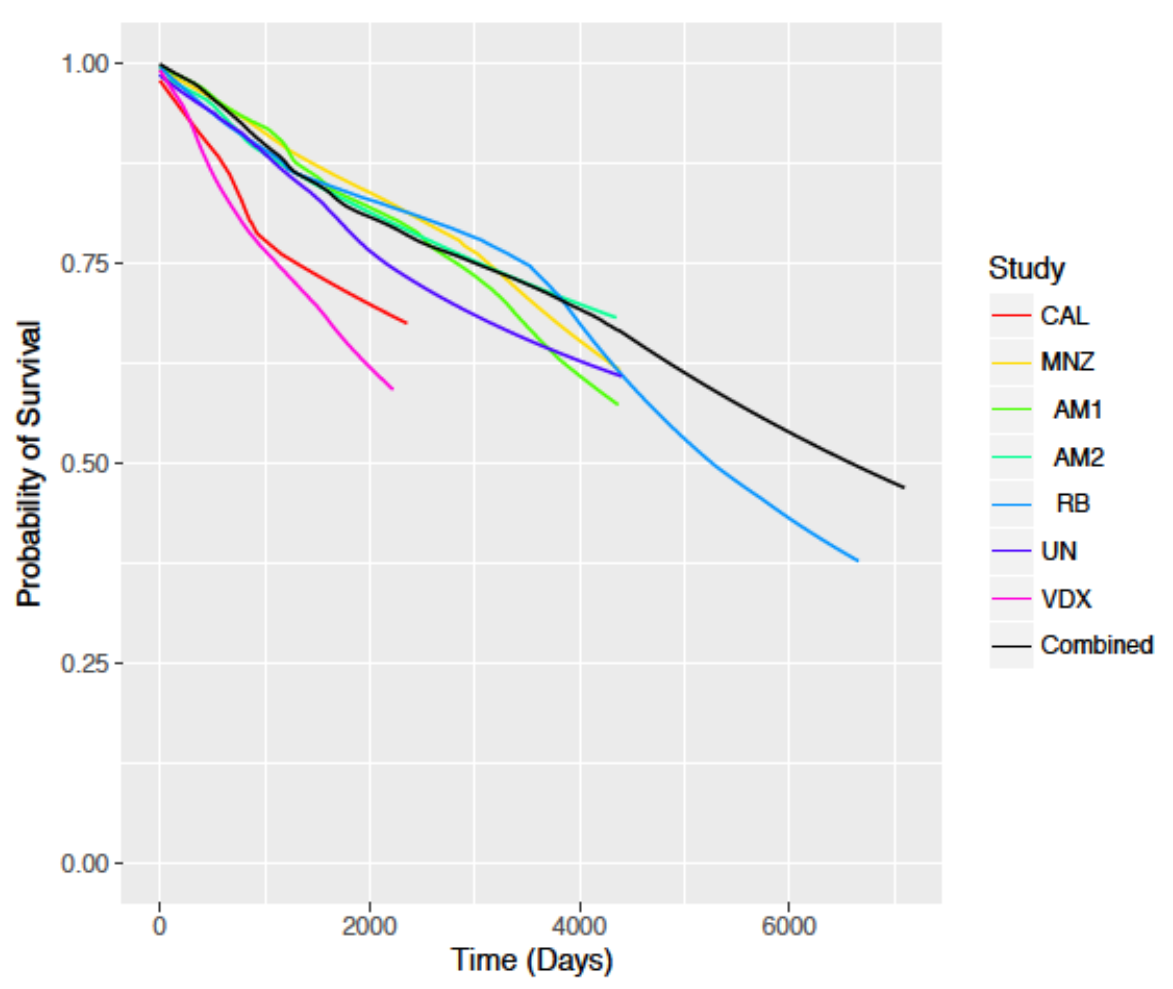
Average probability of survival of each set and the combination of all sets for breast cancer. We compute an expected survival function for each individual in every dataset using the true cumulative hazard and the linear predictor. We then average these survival functions across patients within each dataset. Colored lines represent average survival functions in each original sets. The black line shows the average survival function of all datasets combined. This figure shows the differences in the "true" model from each original study.

Utilizing true survival models that are identical in both baseline hazard and coefficients moderately reduces the difference between within and across-study validation when training Más-o-menos for both breast and ovarian cancer studies. It greatly reduces the within to across-study performance gap when ridge regression is used for both cancer types (Figure 2, “Same models”). In addition, equalizing only the baseline survival functions but not the model coefficients barely reduces the performance gap (Figure 2, “Same hazard”).

### 3.5 Generalization across data types

We now illustrate the results from simulations involving sequencing-based studies. First, we summarized the results of training and validating the prediction models using the TCGA RNA-seq study. Figure 3 and Supplementary Figure 9 show the comparison of results evaluating prediction models across microarray and RNA-seq data types, with those using only the microarray studies. We observed high similarity in the distribution of accuracy estimates between using only microarray studies, and a combination of RNA-seq and microarray studies. Including only highly comparable genes in the original studies brings down the cross-validation performance. Among the sources of heterogeneity we investigate, enforcing identical “true” models is most effective in reducing the performance gap, while maintaining the cross-validation performance at a comparable level to the “baseline” simulations.

These observations are further validated in the metagenomic studies with binary outcome of type-II diabetes vs. control (Figure 2). We observed high variance in AUC scores of models in across-study validation compared to cross-validation. Reducing heterogeneity in clinical factors and in gene measurements barely influences the difference between within to across-study validation. Using identical “true” models greatly reduces the performance gap.

These results demonstrate generalizability of key results across various data types (gene expression microarray, RNA-seq, and whole-metagenome shotgun sequencing), health outcomes (overall survival for ovarian and breast cancers, and type-II diabetes), known sources of heterogeneity (age, tumor grade and size, suboptimal ovarian carcinoma debulking, and BMI), as well as for both time-to-event and binary outcomes.

## 4. Discussion

It is commonly assumed that heterogeneity in experimental platforms or procedures, and differences in patient cohorts, compromise the comparability of independent datasets and the application of omics-based prediction models across studies. This could be addressed by minimizing potential sources of heterogeneity, for example by enforcing precise criteria for patient inclusion. However, such narrowing has costs in sample size and potential generalizability of findings. To the best of our knowledge, no study has carried out a systematic approach to assessing the impact of suspected sources of heterogeneity on the across-study performance of prediction models. We used several compendia of datasets, generated from microarray, RNA sequencing, and whole-metagenome shotgun microbiome sequencing, with study heterogeneity from known and unknown sources, to perform this exploration. We emphasize that the presence of heterogeneity between studies, both measured and unmeasured, is necessary to our investigation.

When training and validating prediction models in these collections of studies, we observed a discrepancy in performance for models validated in fully independent studies when compared to standard cross-validation. For risk prediction of overall survival in compendia of breast and ovarian cancer datasets, independent validation statistics were 0.04 worse on the C-index scale when compared to cross-validation. For predictions of type-II diabetes patients vs. controls from stool metagenomes, accuracy is more than 0.1 worse on the AUC scale in across-study validation compared to within-study validation. These differences in model performance are widespread and sufficiently sizable to question the utility of cross-validation for deciding whether to pursue further development of a prediction model developed and validated on a single dataset. We thus investigated the contributions of known and unknown sources of heterogeneity to this discrepancy. In simulations mimicking these compendia of studies, spanning various data generation technologies and types of outcome, reducing heterogeneity in important clinical covariates did not reduce the discrepancy between CV and CSV. This finding highlights that it should not be assumed that known differences in the composition of different cohorts will negatively impact the application of prediction models across them, or that stricter inclusion criteria will improve the models’ cross-study validation. Several factors are in play. Covariates define strata that are generally associated with different degrees of predictability —in some strata the predictors may be more effective than in others in predicting the outcome. Thus, rebalancing covariates may result in better or worse prediction accuracy depending on which strata are given greater weight. Also, covariate mix in the training sample affects properties of the training algorithm, particularly when relevant covariates are not or cannot be modeled explicitly. Thus, it is difficult to draw general conclusions from individual case studies, but it should not be assumed that stricter inclusion criteria will improve prediction models.

Nonetheless, it is interesting to associate the changes in proportions of clinical covariates to the changes in cross-study validation accuracy. We investigated this association with mixed-effect models in the collections of cancer microarray studies where some covariates strongly affect survival, but found that their marginal distributions do not have much impact on the cross-study stability of predictions of survival. Heterogeneity in the prevalence of covariates like debulking and patient age for ovarian cancer can impact overall survival in different cohorts, but not the ability to predict overall survival.

Similarly, in these compendia of datasets spanning at least 11 different labs and various microarray and sequencing platforms, enforcing good expression measurement comparability through selection of genes with high Integrative Correlation (Parmigiani *and others*, 2004; Garrett-Mayer *and others*, 2008) only moderately closed the gap between cross-study validation and cross-validation in certain data algorithm combinations. Ensuring fully identical models of the association between gene expression and outcome for each study is most powerful in reducing this discrepancy. Thus in these datasets, the most important sources of heterogeneity from the perspective of cross-study validation are likely to be those affecting the relationship between predictors and outcome, and are not likely to be included in the published datasets. For example these could arise from different relationships between covariates and unmeasured confounders, or from different marginal distributions of these confounders.

We used a hybrid parametric/non-parametric bootstrap simulation approach to generate potentially different outcomes for simulated samples that are resampled from the same individual. The parametric step made it possible to simulate the removal of unmeasured confounding through the equalizing of the data-generating models with respect to coefficients and baseline hazards. Theoretically, we could perform rebalancing of covariates in a fully non-parametric approach. However, this would require an extension of the .632 or related approaches (Efron and Tibshirani, 1997) to correct for over-optimism in estimated model performance caused by resampling of *both* individuals within studies and entire studies within the collection. To the best of our knowledge, such a method is not yet described for a cross-study bootstrap. This would be an interesting area for future research.

This study has several limitations. We focus on AUC and C-Index to evaluate the discrimination accuracy of the prediction models, even though these statistics are not directly relevant to clinical implementation. However, they provide advantages of simplicity while adequately capturing the phenomenon of degraded cross-study validation performance relative to cross-validation. The additional model selection in determining thresholds, and necessary assumptions about prevalences to calculate positive and negative predictive values, are complicating factors that may distract from rather than provide additional insight from the basic phenomenon of degradation in cross-study validation performance. Furthermore sampling designs of the studies analyzed are not amenable to validation of positive/negative predictive value without further simulation. In independent validations of predictive value, we would expect prevalence to be an additional source of potential heterogeneity between study populations, but for the conclusions of this manuscript to remain relevant.

We focus mainly on altering one source of heterogeneity at a time. Altering multiple factors yields results that have a less clear interpretation. We report these results in the supplement (Supplementary Figures 9 and 10) to document the interactions between different sources of heterogeneity. We could only analyze clinical strata available in sufficient numbers in these datasets: patient age, tumor size and grade for breast cancer; age and debulking status for ovarian cancer; BMI for type-II diabetes. But our proposed approach shows how the impact of known sources of study heterogeneity can be assessed for their impact on prediction modeling, and that the most obvious heterogeneity may not be the most important.

Despite the limitations, our work has several novel and important contributions. We introduce a novel approach to quantify the impacts of heterogeneity in observed confounders, predictor covariance, and unmeasured confounding on cross-study prediction accuracy. In addition, our results of rebalancing covariates have an important implication - that it is questionable whether studying more clinically homogeneous groups justifies the loss of sample size in practice. This is a common but relatively unexamined practice that we challenge. We developed *simulatorZ* which automates all steps of these simulations including covariate balancing. One powerful feature of *simulatorZ* is that it can simulate data from one or more “omics” studies in a highly realistic way compared to typical synthetic data. Realistic, data-driven, simulated data is essential in evaluating newly developed computational methods. Recent studies have mentioned the utility of *simulatorZ* in evaluating methods for proteomics data (Gatto *and others*, 2015), and in validating replicable cross-study predictors for personalized medicine (Patil and Parmigiani, 2018). By publishing *simulatorZ*, as well as a code repository to reproduce the results of this paper, we hope to encourage further investigation of the effects of study heterogeneity in other predictive modeling contexts.

## 5. Supplementary Material

Supplementary material is available online at http://biostatistics.oxfordjournals.org.

## References

Aalen, Odd. (1978). Nonparametric inference for a family of counting processes. The Annals of Statistics 6(4), 701–726.

Abubucker, Sahar, Segata, Nicola, Goll, Johannes, Schubert, Alyxandria M, Izard, Jacques, Cantarel, Brandi L, Rodriguez-Mueller, Beltran, Zucker, Jeremy, Thiagarajan, Mathangi, Henrissat, Bernard, White, Owen, Kelley, Scott T, Methè, Barbara, Schloss, Patrick D, Gevers, Dirk, Mitreva, Make-donka *and others*. (2012, 13 June). Metabolic reconstruction for metagenomic data and its application to the human microbiome. PLoS computational biology 8(6), e1002358.

Bender, Ralf, Augustin, Thomas and Blettner, Maria. (2005, 15 June). Generating survival times to simulate cox proportional hazards models. Stat. Med. 24(11), 1713–1723.

Bernau, Christoph, Riester, Markus, Boulesteix, Anne-Laure, Parmigiani, Gio-vanni, Huttenhower, Curtis, Waldron, Levi and Trippa, Lorenzo. (2014, 15 June). Cross-study validation for the assessment of prediction algorithms Bioinformatics 30(12), i105–i112.

Binder, Harald and Schumacher, Martin. (2008, 10 January). Allowing for mandatory covariates in boosting estimation of sparse high-dimensional survival models. BMC Bioinformatics 9, 14. CoxBoost.

Castaldi, Peter J, Dahabreh, Issa J and Ioannidis, John P A. (2011, 7 May). An empirical assessment of validation practices for molecular classifiers. Brief. Bioinform. 12(3), 189–202.

Chang, Lo-Bin and Geman, Donald. (2015, November). Tracking Cross-Validated Estimates of Prediction Error as Studies Accumulate. Journal of the American Statistical Association 110(511), 1239–1247.

Cortes, Corinna, Mohri, Mehryar, Riley, Michael and Rostamizadeh, Afshin. (2008). Sample selection bias correction theory. CoRR abs/0805.2775.

Donoho, David and Jin, Jiashun. (2008). Higher criticism thresholding: Optimal feature selection when useful features are rare and weak. Proceedings of the National Academy of Sciences 105(39), 14790–14795.

Efron, Bradley and Tibshirani, Robert. (1997). Improvements on cross-validation: the 632+ bootstrap method. Journal of the American Statistical Association 92(438), 548–560.

Ganzfried, Benjamin Frederick, Riester, Markus, Haibe-Kains, Benjamin, Risch, Thomas, Tyekucheva, Svitlana, Jazic, Ina, Wang, Xin, Victoria, Ahmadifar, Mahnaz, Birrer, Michael J, Parmigiani, Giovanni, Huttenhower, Curtis *and others*. (2013, 2 April). curatedOvarianData: clinically annotated data for the ovarian cancer transcriptome. Database 2013, bat013.

Garrett-Mayer, Elizabeth, Parmigiani, Giovanni, Zhong, Xiaogang, Cope, Leslie and Gabrielson, Edward. (2008, April). Cross-study validation and combined analysis of gene expression microarray data. Biostatistics 9(2), 333–354.

Gatto, Laurent, Hansen, Kasper D, Hoopmann, Michael R, Hermjakob, Henning, Kohlbacher, Oliver and Beyer, Andreas. (2015). Testing and validation of computational methods for mass spectrometry. Journal of proteome research 15(3), 809–814.

Haibe-Kains, Benjamin, Desmedt, Christine, Loi, Sherene, Culhane, Aedin C, Bon-Tempi, Gianluca, Quackenbush, John and Sotiriou, Christos. (2012, 18 January). A three-gene model to robustly identify breast cancer molecular subtypes. J. Natl. Cancer Inst. 104(4), 311–325.

Hartley, H O and Sielken, R L Jr. (1975, 1 June). A “Super-Population viewpoint” for finite population sampling. Biometrics 31(2), 411–422.

Haybittle, JL, Blamey, RW, Elston, CW, Johnson, Jane, Doyle, PJ, Campbell, FC, Nicholson, RI and Griffiths, K. (1982). A prognostic index in primary breast cancer. British journal of cancer 45(3), 361.

Hoerl, Arthur E and Kennard, Robert W. (1970). Ridge regression: Biased estimation for nonorthogonal problems. Technometrics 12(1), 55–67.

Hu, Zhiyuan, Fan, Cheng, Oh, Daniel S, Marron, J S, He, Xiaping, Qaqish, Bahjat F, Livasy, Chad, Carey, Lisa A, Reynolds, Evangeline, Dressler, Lynn, Nobel, Andrew, Parker, Joel, Ewend, Matthew G, Sawyer, Lynda R, Wu, Junyuan, Liu, Yudong, Nanda, Rita, Tretiakova, Maria, Ruiz, Orrico, Alejandra, Dreher, Donna, Palazzo, Juan, P, Perreard, Laurent, Nelson, Edward, Mone, Mary, Hansen, Heidi, Mullins, Michael, Quackenbush, John F, Ellis, Matthew J, Olopade, Olufunmilayo I, Bernard, Philip S *and others*. The molecular portraits of breast tumors are conserved across microarray platforms. BMC Genomics 7, 96.

König, Inke R. (2011, May). Validation in genetic association studies. Brief. Bioinform. 12(3), 253–258.

Leek, Jeffrey T., Scharpf, Robert B., Bravo, Hèctor Corrada, Simcha, David, Langmead, Benjamin, Evan Johnson, W, Geman, Donald, Baggerly, Keith and Irizarry, Rafael A. (2010, 14 September). Tackling the widespread and critical impact of batch effects in high-throughput data. Nat. Rev. Genet. 11(10), 733–739.

Ma, Shuyi, Sung, Jaeyun, Magis, Andrew T, Wang, Yuliang, Geman, Donald and Price, Nathan D. (2014). Measuring the effect of inter-study variability on estimating prediction error. PLoS ONE 9(10), e110840.

Nelson, W. (1969). Hazard plotting for incomplete failure data. Journal of Quality Technology 1, 27–52.

Nelson, W. (1972). Theory and applications of hazard plotting for censored failure data. Technometrics 14, 945–965.

Parmigiani, Giovanni, Garrett-Mayer, Elizabeth S, Anbazhagan, Ramaswamy and Gabrielson, Edward. (2004, 1 May). A cross-study comparison of gene expression studies for the molecular classification of lung cancer. Clin. Cancer Res. 10(9), 2922–2927.

Pasolli, Edoardo, Schiffer, Lucas, Manghi, Paolo, Renson, Audrey, Obenchain, Valerie, Truong, Duy Tin, Beghini, Francesco, Malik, Faizan, Ramos, Marcel, Dowd, Jennifer, B *and others*. (2017). Accessible, curated metagenomic data through experimenthub. Nature methods 14(11), 1023.

Patil, Prasad and Parmigiani, Giovanni. (2018). Training replicable predictors in multiple studies. Proceedings of the National Academy of Sciences 115(11), 2578–2583.

Riester, Markus, Wei, Wei, Waldron, Levi, Culhane, Aedin C, Trippa, Lorenzo, Oliva, Esther, Kim, Sung-Hoon, Michor, Franziska, Huttenhower, Curtis, Parmigiani, Giovanni, *and others*. (2014, May). Risk prediction for late-stage ovarian cancer by meta-analysis of 1525 patient samples. J. Natl. Cancer Inst. 106(5).

Simon, Richard M, Paik, Soonmyung and Hayes, Daniel F. Use of archived specimens in evaluation of prognostic and predictive biomarkers. J. Natl. Cancer Inst. 101(21), 1446–1452.

Uno, Hajime and Inoue, Eisuke. (2017). On estimating predictive performance measures of risk prediction models with external validation data. In: JSM Proceedings. pp. 1156–1161.

Waldron, Levi, Haibe-Kains, Benjamin, Culhane, Aedın C, Riester, Markus, Ding, Jie, Wang, Xin Victoria, Ahmadifar, Mahnaz, Tyekucheva, Svitlana, Bernau, Christoph, Risch, Thomas, Ganzfried, Benjamin Frederick, Huttenhower, Curtis, Birrer, Michael, *and others*. (2014, May). Comparative meta-analysis of prognostic gene signatures for late-stage ovarian cancer. J. Natl. Cancer Inst. 106(5).

Xu, Joshua, Gong, Binsheng, Wu, Leihong, Thakkar, Shraddha, Hong, Huixiao and Tong, Weida. (2016). Comprehensive assessments of rna-seq by the seqc consortium: Fda-led efforts advance precision medicine. Pharmaceutics 8(1), 8.

Zhao, Sihai Dave, Parmigiani, Giovanni, Huttenhower, Curtis and Waldron, Levi. (2014, 23 July). Más-o-menos: a simple sign averaging method for discrimination in genomic data analysis. Bioinformatics.

